# A flexible point and variance estimator to assess bird/bat fatality from carcass searches

**DOI:** 10.1101/2021.08.19.456983

**Authors:** Moritz Mercker

**Affiliations:** Bionum GmbH –Consultants in biological statistics, Hamburg, Germany; Institute of Applied Mathematics (IAM) Heidelberg University, Heidelberg, Germany

**Keywords:** bootstrap, carcass estimator, power line, Monte-Carlo simulation, resampling, wildlife fatality, wind turbine, collision rate

## Abstract

Estimation of bird and bat fatalities due to collision with anthropogenic structures (such as power lines or wind turbines) is an important ecological issue. However, searching for collision victims usually only detects a proportion of the true number of collided individuals. Various mortality estimators have previously been proposed to correct for this incomplete detection, based on regular carcass searches and additional field experiments. However, each estimator implies specific assumptions/restrictions, which may easily be violated in practice. In this study, we extended previous approaches and developed a versatile algorithm to compute point and variance estimates for true carcass numbers. The presented method allows for maximal flexibility in the data structure. Using simulated data, we showed that our point and variance estimators ensured unbiased estimates under various challenging data conditions. The presented method may improve the estimation of true collision numbers, as an important pre-condition for calculating collision rates and evaluating measures to reduce collision risks, and may thus provide a basis for management decisions and/or compensation actions with regard to planned or existing wind turbines and power lines.

## Introduction

Anthropogenic structures, particularly those including thin or fast-moving components, represent a potential collision risk for birds and bats. Prominent examples are wind turbines and power lines [13]. In order to quantify and minimize the related ecological impacts, it is important to be able to estimate the number *N* of fatalities attributable to collision with a certain structure over a specific period of time. However, estimation of *N* based on regular carcass searches is complicated, given that the number of carcasses found, *N*^*F*^, is usually an underestimate of the true number *N*. This is mainly because [20]: (1) not all collided animals die within the search area; (2) not all carcasses persist until the next search (due to decomposition and removal by scavengers); and (3) not all the remaining carcasses are detected by the searchers. The situation is further complicated by the fact that carcasses may be overlooked in one search but detected in a subsequent search (termed ‘bleed-through’ [20]).

One possible way to correct for this underestimation is to perform additional experiments to estimate the corresponding rates of “dying outside”, removal, detection, and decomposition. Such experiments, usually involving appropriate regression models, allow the development of corresponding correction factors that can subsequently be multiplied by raw count numbers, similar to the method of sampling weights [23] in the context of sightability models [14, 24, 30].

Great efforts have been made during the last decade to develop methodologies for estimating *N* using an appropriate estimator 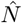 (for overview see for example [3, 20]). However, although various estimators exist, each includes different assumptions and is thus limited to some specific cases. Typical restrictions include the regularity of the search intervals, the underlying distribution function of the carcasses, the constancy or strength of detection/persistence rates, and the number, type, and dependence of the covariates influencing the persistence and detection rates [21, 20]. Furthermore, the estimators have not always taken bleed-through into consideration. As a result of these individual limitations, each existing estimator produces bias under certain conditions [19]. There is thus still a need for a universal estimator that can produce good-quality estimates under general conditions [4, 3]. In addition to providing an appropriate point estimator 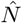, variance estimates (e.g., including confidence intervals (CIs)) are also required. However, during the calculation of 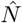 (based on *N*^*F*^), various uncertainties are approximated by empirical mean values, thus inflating the variance of 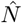. Previous variance estimators have considered at least some of these uncertainties, possibly resulting in an underestimation of CIs.

In this study, we developed a novel universal estimator of wildlife fatalities based on regular carcass searches and additional experiments. Statistically, the estimator consists of the combined application of different regression models to carcass raw count and additional field-experiment data. For variance estimation, these methods were combined with a Monte-Carlo resampling scheme in conjunction with a non-parametric bootstrap method. The presented method allows the calculation of unbiased point and variance estimates for true carcass numbers based on minimal assumptions regarding the raw count and experimental data structures. We verified the functionality of the estimator under various conditions using simulated data, with an appropriate coverage of corresponding CIs. Finally, we developed a procedure for the use of these estimated data within statistical tests.

## Experimental design and derivation of the estimator

### General notation

ℳ represents the total set of carcasses collided with a certain structure in a certain period of time, and *N* = *sum*(ℳ) represents its sum, approximated by the estimator 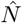 (derived below).

As noted in the Introduction, only a subset ℳ^*F*^ ⊂ℳ of carcasses is found during *J* different regular searches. ℳ^*F*^ is again composed of various single carcasses *M*_*i*_ with the associated covariate vector 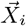, where *i* is a consecutive index. The covariate vector 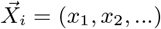 can comprise information about e.g., the date (or search number) and location of carcass discovery, or further covariates related to the detection probability, such as vegetation coverage (c.f., following sections). As mentioned above, additional field experiments are necessary to estimate the persistence probability *s*(), searcher efficiency *f* (), decomposition time *t*_*d*_(), and the average number of collided birds/bats falling into the search area, *A*_*in*_(), all of which enter the final estimator. These approximations are given in terms of appropriate statistical models *ŝ*(), 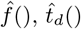, and *Â* _*in*_(), respectively.

We define these models below, and subsequently derive the estimator 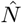, as the sum of pointwise estimates 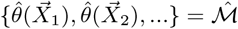.

### Proper and improper uses of 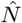

Importantly, 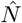 is only valid for estimating carcass numbers *N* of animals that collided with the specific structure/component located within the search area, and for collisions occurring during the investigated time period. In contrast, 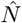 cannot be used for temporal or spatial extrapolation, e.g., to estimate numbers of collisions with structures located outside the search area or occurring before/after the period of time comprising the regular searches. This restriction is based on the fact that bird or bat migration/appearance may differ between different sites or periods [25, 22]. Errors associated with extrapolation therefore cannot be controlled.

However, the potential search area or time interval may be too large to allow regular searches across the entire area or period of time. In this case, the presented approach should be embedded into an appropriate sampling design (e.g., a stratified design [30, 23]), rather than extrapolating values. It is also possible that some sites within the search area cannot be monitored at some time points (e.g., due to flooding events or agricultural use). However, this situation is qualitatively different, because corresponding missing values can be interpolated from existing data, e.g., using the methods of multiple imputation [36].

Especially in the context of impact studies, the average difference in carcass numbers per search between two different sites/structures may be the issue of interest, rather than the total sum *N* [32]. In the simplest form, such a comparison can be made via a *t*-test [16]. However, neither raw counts nor pointwise estimates provide a valid data basis for such a test, and the spread of pointwise estimates among themselves, as well as the individual accuracy of the estimates, should be taken into account. The corresponding statistical methods are presented below (“Carcass estimates within statistical tests”).

### Persistence probability

Persistence probability is frequently assessed experimentally by laying out a number of fresh carcasses within an appropriate area and observing their persistence over at least several days. This allows an estimation of the time of removal after exposure for each individual carcass. Carcasses can be checked every day or more frequently (e.g., using camera traps [26]), given by a scaling factor *α*_*t*_ relative to the number of days. E.g., *α*_*t*_ = 24 indicates that data were collected hourly. Time spans (e.g., denoted by *D*_(*k,l*)_) are given below in units of *α*_*t*_ · *days*.

To use a general model for such repeated measurements over time, we used a regression framework of “survival analysis”, which allowed consideration of an arbitrary additional number and type of covariates [6, 7, 20], and in which not only exponential decay, but various types of decay functions could be compared. The best model (with respect to decay function and covariate combination), *ŝ* (), could then be selected based on the Akaike’s Information Criterion (AIC) [1]. Finally, all underlying model assumptions need to be proven before using *ŝ* () within a corresponding correction factor.

*ŝ*() thus finally depends on a time point *a* (measured from the time of carcass exposure) and an additional number and type of covariates 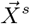. Hence, 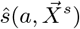 approximates the probability *P*^*pers*^ that a carcass (with a set of certain covariates) still remains after the time interval [0, *a*], thus 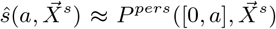. In the following, we were also interested in the persistence probability for later time intervals [*a, b*], i.e., with *a >* 0. Numerous previous studies have approximated this probability via *P*^*pers*^([*a, b*]) ≈ *P*^*pers*^([0, *b* − *a*]) [3, 19]; however, this relationship is only valid if we assume that *P*^*pers*^([*a, b*]) decreases exponentially over time, i.e., has a constant removal rate. This assumption is easily violated, e.g., if the removal rate depends on carcass age. Non-constant removal rates thus produce bias using previous estimators [19].

In the present study, we corrected for this bias approximating *P*^*pers*^([*a, b*]) via

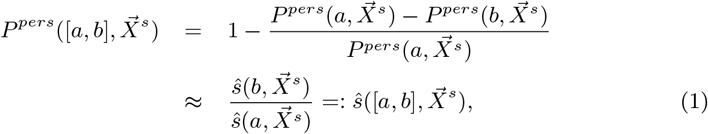

setting 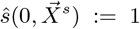 (a carcass always persists in the moment of exposure/falling down). We thus directly calculated the relative change in *P*^*pers*^ between time points *a* and *b*, which is not necessarily equal to the change between 0 and (*b* −*a*) as assumed in previous models. Notably, the right hand side of equation (1) reduces (only) in the exponential case to 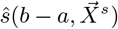 and is thus a generalization of previous approaches.

### Searcher efficiency

Searcher efficiency *f* () can be assessed experimentally by performing “artificial carcass searches”, where the true number and location of carcasses is known to persons other than the searchers. Here, we fitted a generalized linear mixed model (GLMM) [8] (in particular a logistic GLMM) to the data, modelling the binary outcome “detected/not detected” and allowing for an arbitrary number and type of covariates influencing *f* (). Individual observers are usually treated as a random effect to avoid bias connected to stochasticity of the experimental data [20].

We also tested various possible covariates influencing *f* () simultaneously, and subsequently selected the best regression model 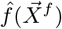 (with appropriate covariates 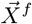) based on the AIC value [1]. As for *ŝ* (), all assumptions necessary for a generalization (e.g., described by [16]) should be proven before using 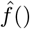 in the context of a correction factor. 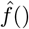 thus represents the probability that a carcass with a certain set of covariates 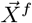 is detected during a search.

### Decomposition time

The decomposition time *t*_*d*_() represents the number of days until a carcass is undiscoverable by a searcher due to its progressive decomposition. Under certain circumstances, *t*_*d*_() can be estimated together with persistence probability *s*(), leading to a more general persistence probability 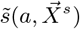, representing the probability that a carcass with certain covariates 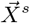 is neither removed nor decomposed after time *a*. However, the time scale and/or temporal dependency may differ between both processes, such that separate experiments assessing *t*_*d*_() are recommended. One possible experimental design would involve the long-term exposure of carcasses protected by cages prior to removal.

However, due to differences in the time scales corresponding to removal and decay, decomposition time often plays a subordinate role compared with removal by scavengers, and the experimental effort should thus be adapted accordingly. Thus, considering the empirical mean 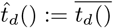 as the simplest possible regression model may be sufficient in many cases, though more sophisticated regression models and additional covariates 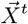 may also be used.

### Proportion of carcasses falling into the search area

It is important to estimate and consider the proportion of collided birds/bats that falls into the search area, *A*_*in*_(), which may be only a minority of individuals under some circumstances [2]. However, experimental estimation of *A*_*in*_() is difficult, because it would also require estimating the number of carcasses within the unsearched area, which is an arbitrarily large area. Furthermore, attributing carcasses to the structure of interest becomes more difficult with increasing distance from the structure. Nevertheless, there are some direct and indirect methods that can be used to experimentally assess *A*_*in*_().

1. If detected carcass numbers within the search area are sufficiently high, a model density function *g*(*x*) can be fitted to the data, representing the relative decay in carcass numbers with increasing distance *x* from the structure. In some cases, a normal density function may be appropriate (e.g., considering collision with power lines), while in other cases (e.g., if additional ballistic effects of wind turbines are taken into account), more sophisticated models are recommended [18]. Before fitting *g*() to the data, the latter should be corrected for persistence probability, searcher efficiency, and decomposition time. Finally, an integration of *g*() over the entire area compared with the integral truncated to the search area leads to the desired estimate *Â*_*in*_(). Ideally, this is carried out separately for the data for each search (or, because the data are usually too sparse, for data pooled over several longer time intervals), in order to also assess the variance of *A*_*in*_(). However, this approach may underestimate the number of injured birds flying longer distances before dying from the consequences of the collision (known as “crippling bias” [2]).
2. Regular and systematic visual observations of flying animals within the immediate surroundings of the structure (including direct observation of collisions) may help to estimate the number of collided birds that do not fall into the search area. However, this approach is associated with several potential problems: such observations are biased in favour of larger birds, and it is almost impossible to decide if a collided bird that continues to fly has been mortally injured. Finally, direct observations of collisions are rare and thus require a relatively large experimental effort to obtain meaningful numbers.
3. The use of radio-tagged animals and molecular tracking methods / forensic approaches has also been used/proposed to improve estimates of crippling bias [2, 5]. However, these methods can only be applied to relatively small numbers of animals and involve extensive experimental effort.

### Deducing the estimator

Be 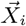 the covariate vector associated with a certain carcass with index *i*, found at search number *j* ∈ {1, 2, …, *J*}. 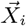 should include all covariates appearing in the “best regression models” 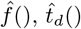, and *Â*_*in*_() (c.f., previous sections). Furthermore, *D*_(*j,k*)_ represents the number of days between the current search and search number *k*, scaled by the factor *α*_*t*_ (c.f., section “Persistence probability”). In the following, we developed a correction term 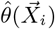, approximating, for each detected carcass *i*, the total number of similar carcasses (i.e., with similar covariate values) that collided with the structure of interest, correcting for carcasses that had been removed by scavengers, overlooked, fallen outside the search area, or decomposed.

The formula for 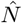 estimating the true carcass number *N* was based on the combined application of different regression models applied to raw counts and additional experimental data. 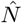 was defined as the sum of corrected raw counts 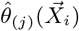 and possibly additional imputed missing data 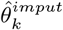 by

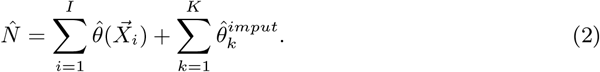

Especially, each value 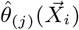 is given by

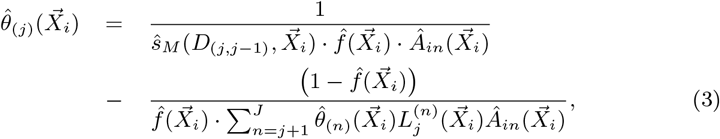

where the index *j* indicates that the carcass was found during the *j*-th search. This equation has to be solved iteratively to obtain 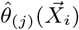 : Starting with 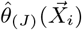 (which is well defined, c.f. Supporting Information), 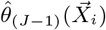 is uniquely determined, which again yields 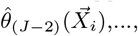, up to 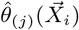.

Furthermore, within Eq. 3 we need the following definitions:

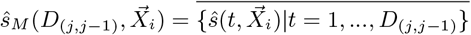

and

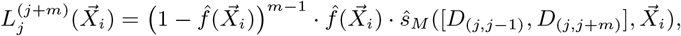

which are again based on Eq. 1. Please see the Supporting Information for more details on the successive derivation of this formula.

## Variance estimation

### General approach

Calculating the variance of 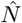 is challenging, and analytical methods quickly reach their limits, especially if the models include a high degree of complexity [27]. Available analytical estimators (e.g., Ref. [30]) consider much simpler sightability models compared with the present model. Some parts of these estimators have been shown to be biased, and additional implementation errors in standard evaluation software have increased the variance bias in several publications [14].

The current study applied several different correction terms in a nonlinear manner, allowing the use of a bootstrap method as a possible alternative to an analytical estimation of CIs [23, 27, 15]. We thus developed a corresponding resampling scheme based on Monte-Carlo-simulations in conjunction with a non-parametric bootstrap method.

### Factors inflating the variance of 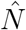

During calculation of the estimate 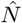, various uncertainties/random processes are approximated by empirical means, thus inflating the final variance:

1. a carcass may or may not fall into the search area;
2. a carcass may or may not be removed by scavengers;
3. the exact time point of dying is unknown for a carcass, leading to additional uncertainties regarding the persistence probability;
4. a remaining carcass may or may not be detected by a searcher (after one or more searches);
5. the predicted values of the statistical models entering the estimator (and deduced from additional experiments) are associated with uncertainties;
6. imputation of missing values is also associated with uncertainties.

All these points are incorporated within the following bootstrap algorithm and inflate the variance estimate.

### Resampling scheme

We were interested in the average spread of 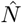 around the true value *N*. Because only one realization of 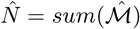 was available, a corresponding bootstrap estimate was based on resamples of 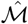, virtually imitating and integrating all six above-mentioned sources of uncertainty. For each resample, this leads to a newly generated “virtual set of detected carcasses” 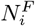, from which the estimate 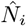 can be calculated. Repeating these steps *n*_*boot*_ times produces a set of estimates 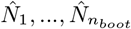 spread around 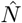, imitating the spread of 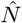 around the true value *N*. Variance bias due to 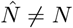 (non-pivotality) and corresponding corrections is considered below. Please see the Supporting Information for more details regarding this non-standard resampling scheme.

### Calculation of CIs and bias correction

The 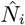 values appeared to be significantly non-normally distributed in all our studies (e.g., induced by the fact that 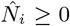 holds), and standard non-parametric bootstrap methods would thus lead to bias, since 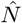 is a non-pivotal quantity: the standard error of 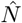 varies with the size of the estimate and it usually holds that 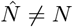. Appropriate correction procedures should thus be applied to prevent poor coverage probability. To account for both the non-normality of the data and non-pivotality, we applied non-parametric bias-corrected and accelerated (BCa) bootstrap methods [12] to calculate CIs based on the population of 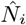 values.

## Computation and performance

### Software and packages

The estimator 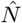 and corresponding CIs were calculated using the open-source software **R** version 3.1.0 [29]. We used the packages **survival** [33] for survival regression in the context of persistence probability, **mgcv** [35] for the GLMMs and generalized additive models (the latter realizing data imputation), and **ggplot2** [34] for visualizations. BCa CIs were calculated using the package **simpleboot** [28].

### Simulation studies

To prove the functionality of the developed point and variance estimators, we tested our algorithms using simulated data, with a known true number of collided individuals *N*. We investigated estimator bias under various conditions, as well as the coverage probability depending on sample size. We used virtual data reflecting highly heterogeneous/difficult data conditions.

#### Virtual carcass search data

We prescribed a search area divided into two sites, and a highly variable mortality rate (on average increasing) over time and between the two sites. Furthermore, if not stated otherwise, we assumed that 10 different searches were conducted, with time intervals between searches of 1 to 4 days. Covariate values for each individual carcass were generated randomly leading to persistence and detection rates ranging between 0.2 and 0.8.

#### Virtual field experiments

We also created artificial experimental data to calculate *ŝ*() 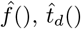, and *Â*_*in*_(), as described above. Here, 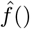 showed non-constant dependence on the covariate, *ŝ*() decayed non-exponentially over time (i.e., having a highly variable removal rate) with a half-life of 6 days, *t*_*d*_() was set to 10 days, and *Â*_*in*_() was based on randomly generated values from the normal distribution *N* (*mean* = 0.8, *sd* = 0.1). For the sake of simplicity, the pseudo-experimental data did not consider individual searchers or carcasses, and standard logistic regression models (generalized linear models) rather than GLMMs were therefore applied.

### Point estimator performance under various conditions

In order to validate the performance of the estimator under non-standard conditions, we created and investigated different challenging/non-standard datasets by modifying the above simulated data, always considering *N* = 20 carcasses (for the specific modifications/challenges, c.f., Fig. 1). We repeated the following steps 10.000 times for each modified set of data:

1. allocation of a random covariate value and a random time point of dying (between two prescribed searches) for each individual carcass;
2. randomly replacing the data of one date/site combination by missing values (resulting in 5% missing values);
3. performing “virtual detection/removal/falling outside” using Monte-Carlo methods based on *ŝ*(), 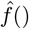, and *Â*_*in*_() applied to corresponding covariates and persistence times. Here, we used resamples of experimental data reflecting the fact that the true functions were unknown. If a carcass was not removed or detected, or fell outside or exceeded the decomposition time given by a resample of *t*_*d*_(), it was shifted to the following search;
4. based on the resulting virtual raw count data, 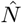 was estimated as described above.

**Figure 1:**
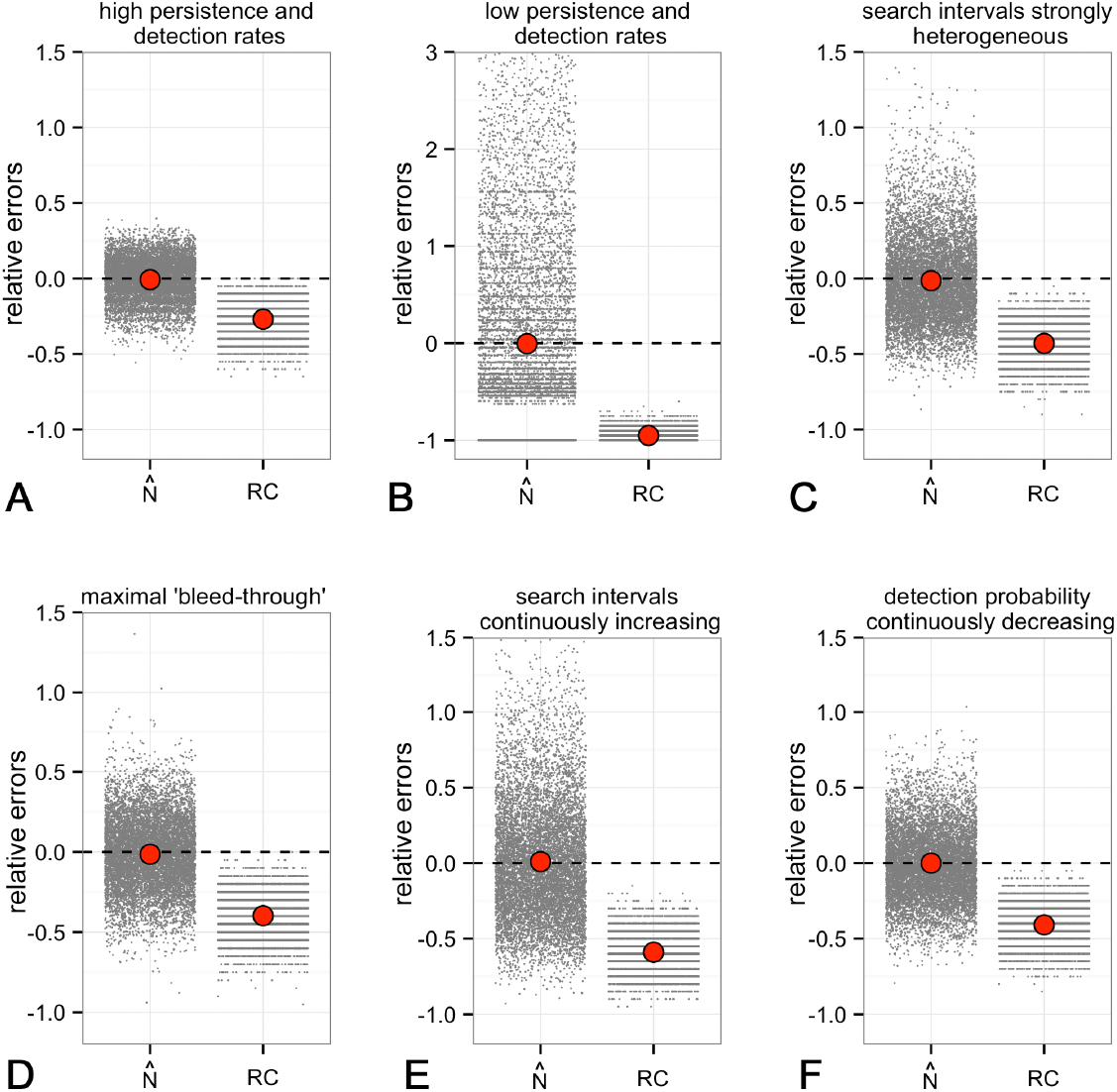
Relative bias of the estimator 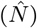 and respective raw counts (*RC*) (both compared to the true number *N*) of simulated datasets under various challenging data conditions. Small grey dots: single data values (randomly scattered in x-direction); large red dots: empirical mean values; dashed line: zero bias. Detailed parameters: (A)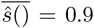, search interval 1 day. (B) 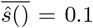, search interval 14 days. (C) Search intervals varied between 1 and 20 days. (D) Maximal ‘bleedthrough’ due to 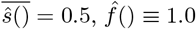. (E) Search intervals increased continuously from 1 to 20 days. (F) 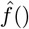 decreased continuously from 0.9 to 0.1.

Thus, for each dataset we created a population of estimates that could be examined with regard to bias, considering the relative errors of the estimate, 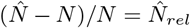 compared with those of the corresponding raw count, 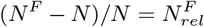. All results are shown in Figure 1. Here, relative errors associated with single estimates (small grey points) were scattered randomly along the x-axis to increase their visibility. Respective empirical mean values of 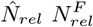 are shown as large red dots.

We first investigated the performance under favourable conditions, i.e., a high (non-constant) detection rate with mean value 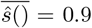 and constant search intervals of 1 day. In this case, the estimates 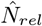 were scattered relatively closely and symmetrically around *N* (Fig. 1 A). In contrast, 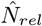 values were spread widely and non-symmetrically around zero-bias if we prescribed very poor detection and persistence conditions setting 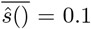 and search intervals of 14 days (Fig. 1 B). In contrast to other presented simulation studies, some of the raw count data did not contain any carcasses, leading to a population of 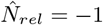 values within the corresponding figure. Furthermore, this example demonstrated that, especially for such poor conditions, 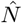 was strongly non-normally distributed around *N*, suggesting non-parametric methods of estimating CIs. In the dataset corresponding to Figure 1, we strongly varied the persistence probabilities by randomly generating search intervals between 1 and 20 days. Here, as in the following examples, 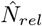 values were relatively symmetrically distributed. In the dataset corresponding to Figure 1 D, we provoked a maximal rate of carcasses found at later searches (‘bleed-through’) by setting 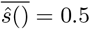 and 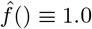. Finally, we continuously increased the search interval from 1 to 20 days (c.f., Fig. 1 E) and continuously decreased the detection probability 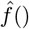 from 0.9 to 0.1 (c.f., Fig. 1 F).

All these conditions showed qualitative and quantitative differences regarding the spread of single estimates around a relative error of zero, but all appeared to be unbiased. In contrast, raw count numbers always underestimated the true number *N*, as expected.

### Assessing coverage probability

We investigated the experimental coverage probability of the calculated CIs, i.e., we verified if the predicted CIs contained in theory the true value *N* with the prescribed nominal value of 0.95. Based on the heterogeneous virtual data introduced above, we generated 11 different datasets comprising *N* carcasses ranging between *N* = 20 and *N* = 70. From each of these sets, we first generated 200 different virtual sets of field data 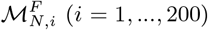 by simulating all different random processes (removal, detection, falling outside,…) based on Monte-Carlo simulations. Based on each set 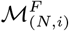, we subsequently calculated 95% CIs based on *n*_*boot*_ = 200 resamples, as described above. Finally, we determined if the calculated CIs contained the true value *N*.

The results are shown in Figure 2. The experimental coverage (black dots) deviated from the nominal coverage, especially for small numbers of *N* (coverage of about 0.91 for *N* ≈ 20), but distinctly converged towards the nominal value for higher values of *N* : Regression analysis of type *f* (*N*) = *b*_0_ + *b*_1_*/N* (red line in Fig. 2) yielded highly significant values *b*_0_ = 5.41 and *b*_1_ = 86.36. This observation fit well with the previous studies of Ref. [11, 10], which showed that BCa intervals could significantly exceed the nominal error rate when the sample size was of the order of 10, but converged towards the nominal rate for larger samples.

**Figure 2:**
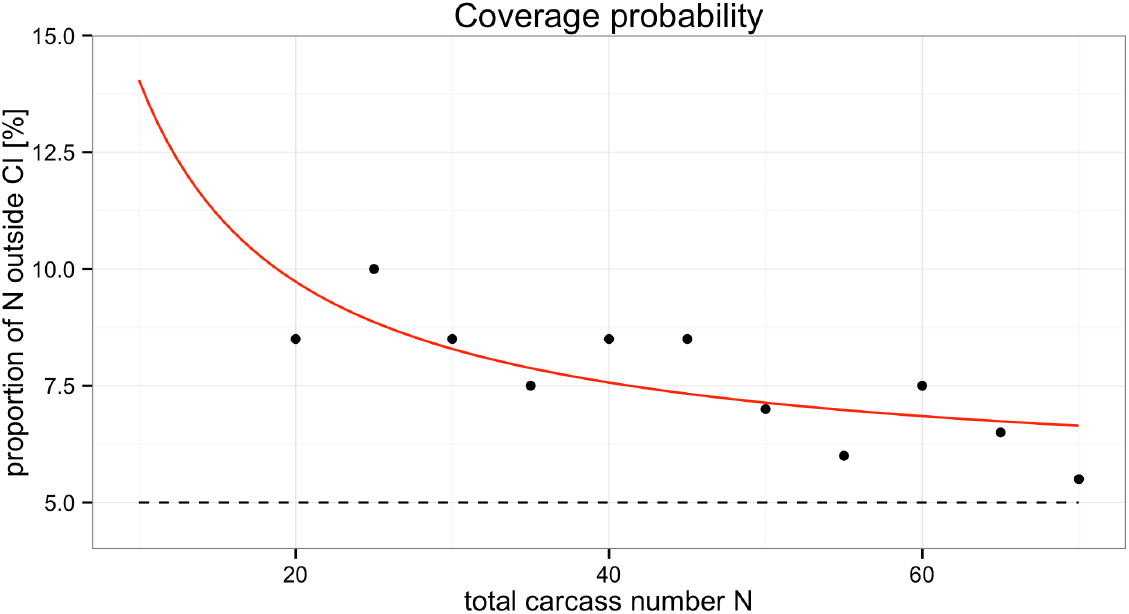
Assessing the true coverage probability for the presented point and variance estimation method. For various simulated datasets *M* containing different total carcass numbers *N* we generated 200 different virtual sets of field data 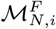 using Monte-Carlo simulations for each dataset (*i* = 1, …, 200). Based on each set 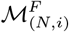, we subsequently calculated 95% CIs based on *n*_*boot*_ = 200 resamples and finally checked if the CIs contained the true value *N*. Experimental coverage values are denoted by black dots; red line indicates the regression curve *f* (*N*) = 5.41 + 86.36*/N*.

The small difference between the experimentally approached value *b*_0_ = 5.41 and the nominal value of 5.0 could be due to the relatively small resample size of *n*_*boot*_ = 200 used in our simulation study. In contrast, resample sizes of *n*_*boot*_ = 2000 are recommended when CIs are bootstrapped [16]. However, assessing coverage probabilities for much larger sample or resample sizes is beyond the scope of the present study, given that 400 estimates each with 200 resamples already resulted, especially for higher values of *N*, in computation times of more than 1 week on a single core, using the virtual data presented above, due to the computationally intense iterative nature of the 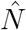 calculations.

### Carcass estimates within statistical tests

Especially in the context of impact studies, the total number of carcasses *N* of animals that collided with a certain structure within a certain period may not be the matter of interest. In contrast, the average difference in carcass numbers between two sites/structure types *A* and *B* can be considered, and may be assessed by performing various simultaneous searches within these two sites, leading to a set of carcass numbers 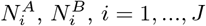. The question is thus if the “means of the carcass-producing processes” [17, 32] differ between these two sites (the latter represented by the two sets 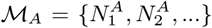 and 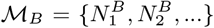. Appropriate statistical tests can be applied to address such questions, ranging from simple *t*-tests to more sophisticated ANOVA- or regression-based before-after control-impact designs [32, 31]. We describe below how to apply statistical tests appropriately to such estimates by propagating all uncertainties to final *p*-values/standard errors, as an approach rarely applied in previous studies.

*Q* is a scalar empirical statistic (e.g., mean or regression coefficient) that should be calculated based on a set of carcasses ℳ to estimate the “true underlying expected value” *Q*^*real*^. Here, we assumed that all carcasses were found, and *Q* was thus one possible realization dispersing around *Q*^*real*^ (c.f., Ref. [32]). In order to assess information about the range possibly comprising *Q*^*real*^, we approximated the variance of *Q*^*real*^ using the sample size and variance of *Q*, the latter based on the dispersion of the individual values ℳ = {*M*_1_, *M*_2_, …}, termed “extrinsic variance”. This is the normal practice by which CIs are calculated.

However, in the current case, additional complexity was introduced because *ℳ* was itself unknown, and our analysis was thus based on the set 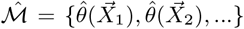, comprising estimates instead of sharp values. This issue is frequently neglected and these estimates are treated as sharp values. However, these uncertainties (“intrinsic variance”) should be added to the extrinsic variance to obtain an unbiased estimate of *Q*.

This can easily be realized within the presented bootstrap framework: as described above, we can generate several Monte-Carlo-based resamples 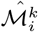 (*i* = 1, …, *n*_*boot*_ and *k* = *A, B*), reflecting (only) uncertainties appearing during estimation (the intrinsic variance). To additionally incorporate the extrinsic variance (reflecting the spread of *Q* around *Q*^*real*^), we subsequently applied the classical bootstrap approach to each of the above resamples and created a resample, termed as 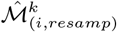, from each 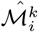. In the simplest case, this would be a common random resample with replacement. However, because count data were considered and additional temporal and/or spatial correlation may exist, more sophisticated resampling models could be appropriate, e.g., including the identification of relevant strata for resampling [9]. Finally, we applied the statistic *Q* to all these “double-resamples”, leading to two classes of resamples 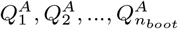 and 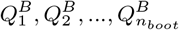, which can be further assessed using appropriate statistical tests (such as a *t*-test or regression analysis).

## Summary and discussion

The presented approach offers a flexible and universal method for assessing point and variance estimates for true carcass numbers resulting from collisions with anthropogenic structures, based on regular carcass searches, the latter being prone to observation bias (not all carcasses are usually detected). In contrast to previous approaches, this novel method does not rely on specific assumptions e.g., regarding the nature of detection/persistence rates, search interval, covariates influencing detection/removal rates, or a specific probability distribution for detected carcasses. Using simulated data, we showed that our estimator was unbiased, even under extreme data situations, e.g., with respect to strongly inhomogeneous search intervals or non-constant detection/removal probabilities. However, a possible shortcoming of this novel method could be its relatively high computational effort: the presented equations contained different iterative elements, meaning that the computation time increased exponentially in line with the number of carcass searches. However, our R-implementation may not have been programmed in the optimal time-efficient manner due to a lack of corresponding skills, and efficient programming in conjunction with a parallelized code could greatly reduce the required computation times. Furthermore, the iterative nature and high complexity meant that the estimator could not be presented in the form of a “easy-to-implement one line-formula”, as for previous approaches [3, 20].

Standardized and unbiased statistical methods are an important prerequisite for estimating and comparing the ecological impacts of different anthropogenic structures (or locations) on bird and bat fatalities, e.g., by guaranteeing comparability. The present model may provide a basis for this process, by offering a universal and unbiased method for estimating point and variance estimates of carcass numbers. Furthermore, the presented bootstrap approach can be easily embedded within various statistical tests, thus allowing the appropriate use of carcass estimates in the context of impact studies.

## Appendix

### 1. Notation

**Table 1:**
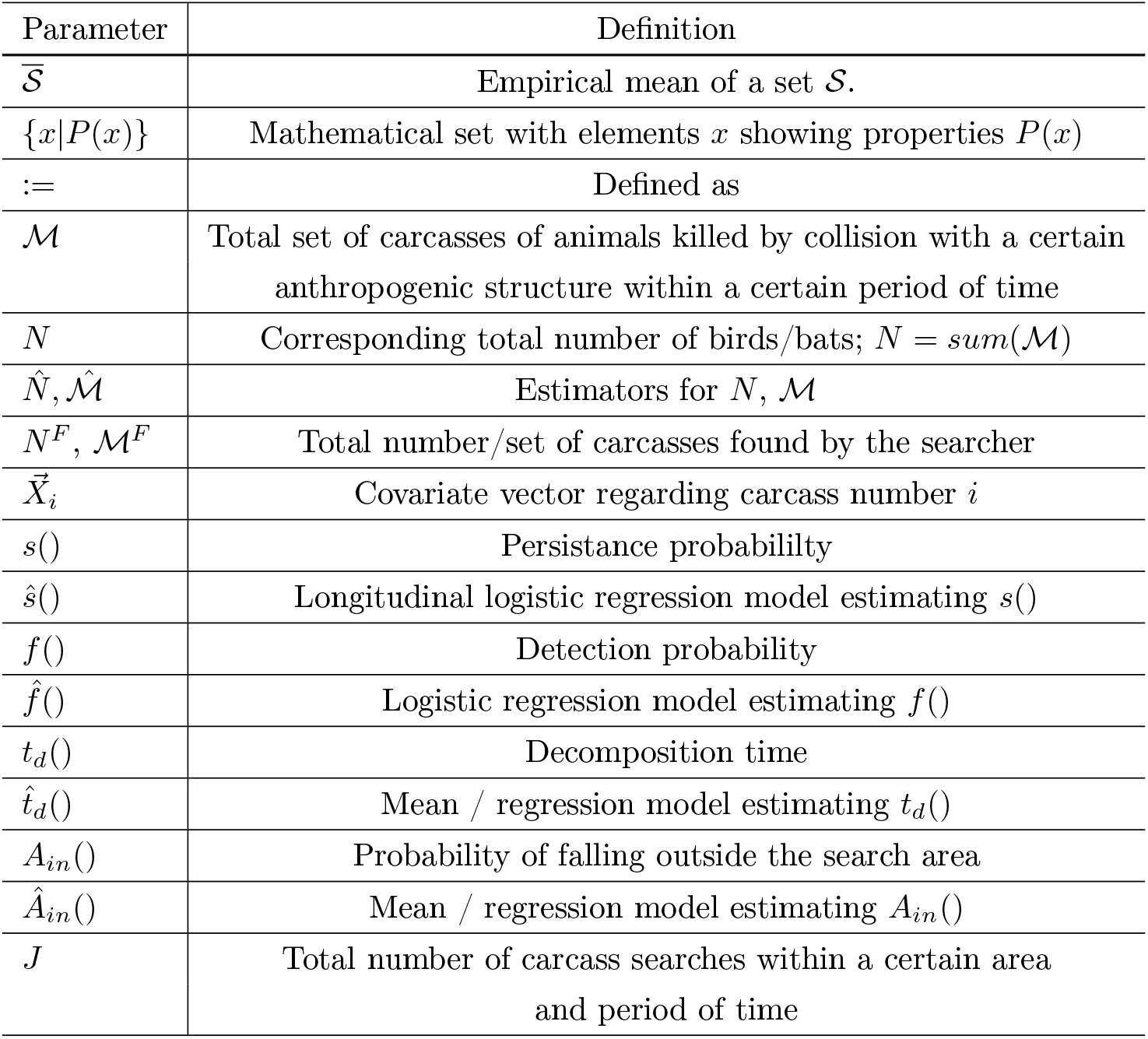
Definitions and notation part I

**Table 2:**
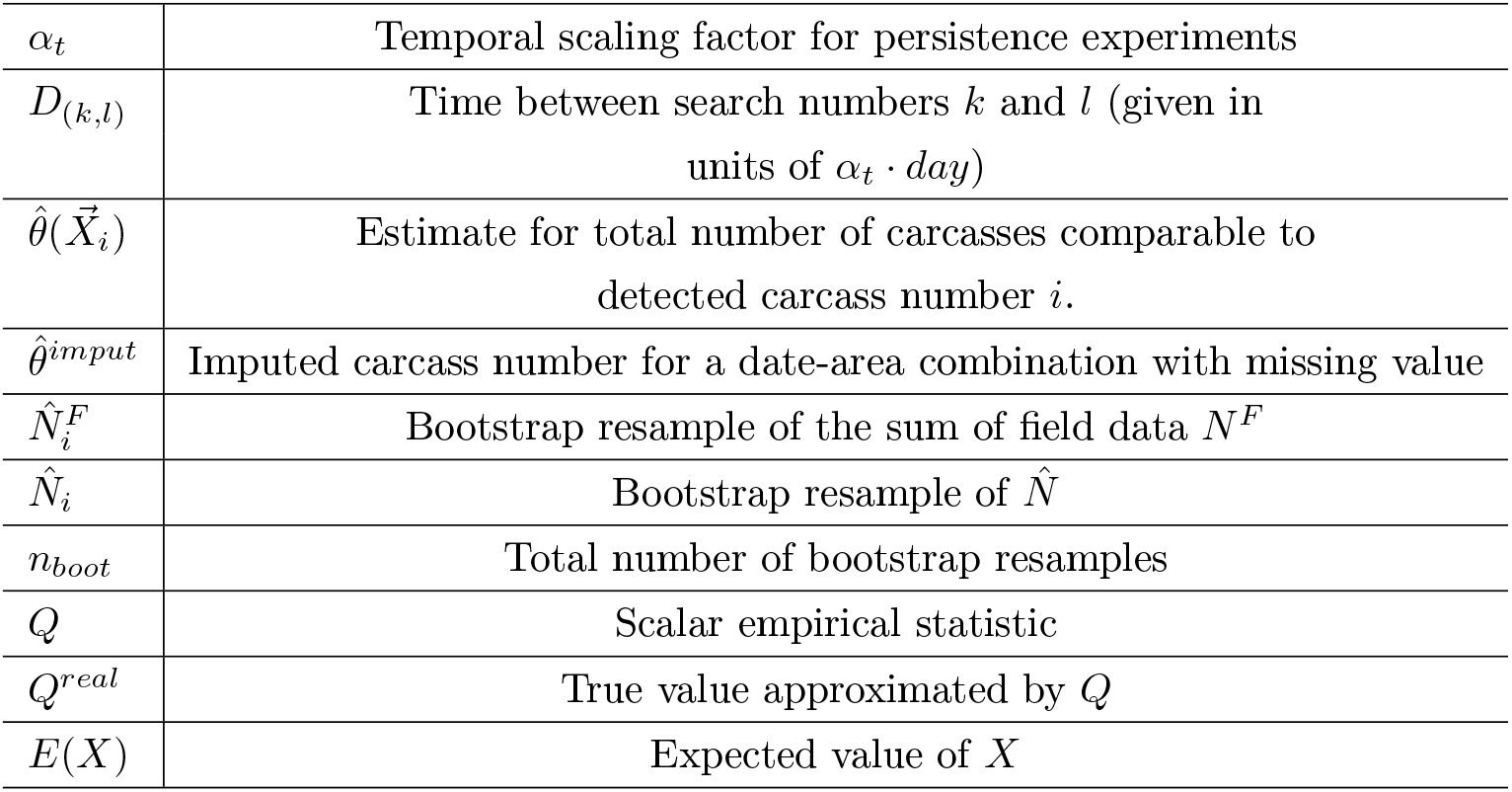
Definitions and notation part II

