## Supporting Information for "A flexible point and variance estimator to assess bird/bat fatality from carcass searches"

Moritz Mercker

### Deducing the estimator

In the following, we present a detailed derivation of the estimator $\hat{N}$ given by Eq. 2 in the main manuscript.

#### Persistence and searcher efficiency

In order to firstly correct the number of carcasses for persistence and searcher efficiency via correction terms $\hat{\theta}^{(s,f)}(\vec{X}_i)$, we extend the "sightability models" as presented in Ref. (1, 3–5). Here, a first naive estimate of the corrected number is given by

$$\hat{\theta}^{(s,f)}_{(j)}(\vec{X}_i) = \frac{1}{\hat{s}_M(D_{(j,j-1)}, \vec{X}_i) \cdot \hat{f}(\vec{X}_i)}. \tag{1}$$

Since the exact time point of dying is unknown for the carcass of interest, $\hat{s}_M(D_{(j,j-1)}, \vec{X}_i)$ is defined as the corresponding empirical mean over all possible time points, hence

$$\hat{s}_M(D_{(j,j-1)}, \vec{X}_i) := \overline{\{\hat{s}(t, \vec{X}_i) | t = 1, ..., D_{(j,j-1)}\}}. \tag{2}$$

Here, we assume that collision probability is equally distributed in time. If collision probability rather shows some typical time-dependent pattern, a corresponding weighting function (depending on daytime) should be multiplied with values $\hat{s}_M(t, \vec{X}_i)$.

Furthermore, it is important to notice that for the first search of the monitoring period, carcasses remaining from unknown earlier time points would bias the estimate. Thus, it is recommended to perform an outstanding intensive carcass search $D_{bef}$ days before the first regular search (without recording these found carcasses), subsequently setting $D_{(1,0)} = D_{bef}$ for the first regular search.

#### Bleed-through

Secondly, we have to incorporate the fact that carcasses can be overlooked in one search, but detected in one of the following searches ("bleed-through"): From the above mentioned $1/\big(\hat{s}_M(D_{(j,j-1)},\vec{X}_i)\cdot\hat{f}(\vec{X}_i)\big) =: C$ carcasses, in average,

$$C\cdot\hat{s}_M(D_{(j,j-1)},\vec{X}_i)\cdot(1-\hat{f}(\vec{X}_i)) = \frac{(1-\hat{f}(\vec{X}_i))}{\hat{f}(\vec{X}_i)} =: K \qquad (3)$$

carcasses remain after the $j$-th search in the field. Thus, $K$ is the amount of carcasses which has neither been removed by scavengers nor detected by searchers until the end of the $j$-th search. Be $D_{(j,j+m)}$ now the number of days until the $(j+m)$-th search after the $j$-th search. In this case, the average probability of finding one of the $K$ carcasses at search number $(j+m)$ is given by

$$\big(1-\hat{f}(\vec{X}_i)\big)^{m-1}\cdot\hat{f}(\vec{X}_i)\cdot\hat{s}_M([D_{(j,j-1)},D_{(j,j+m)}],\vec{X}_i) =: L_j^{(j+m)}, \qquad (4)$$

where

$$\hat{s}_M([D_{(j,j-1)},D_{(j,j+m)}],\vec{X}_i) := \overline{\{\hat{s}([t,D_{(j,j+m)}],\vec{X}_i)|t=1,...,D_{(j,j-1)}\}}. \qquad (5)$$

(for calculation of the $\hat{s}([t,D_{(j,j+m)}],\vec{X}_i)$, c.f., equation (1) main manuscript). Thus, it must hold that one of the $K$ carcasses is found during the $(j+m)$-th search without being found by searchers or scavengers before. However, if one of these $K$ carcasses is found, also a corresponding correction term $\hat{\theta}^{(s,f)}_{(j+m)}$ is applied and biases the estimate. Thus, considering all following searches, alltogether

$$K\cdot\sum_{n=m+1}^{J}\hat{\theta}^{(s,f)}_{(n)}L_j^{(n)} \qquad (6)$$

carcasses "too much" will appear within the estimate $\hat{N}$. Hence, we define the corrected estimate by

$$\hat{\theta}_{(j)}^{(s,f)} = \frac{1}{\hat{s}_M(D_{(j,j-1)}, \vec{X}_i) \cdot \hat{f}(\vec{X}_i)} - K \cdot \sum_{n=j+1}^{J} \hat{\theta}_{(n)}^{(s,f)} L_j^{(n)}. \tag{7}$$

Here, the $\hat{\theta}_{(n)}^{(s,f)}$ values can be determined iteratively: Based on $\hat{\theta}_{(J)}^{(s,f)} = 1/\big(\hat{f}(\vec{X}_i) \cdot \hat{s}_M(D_{(J,J-1)}, \vec{X}_i)\big)$ (i.e., there are no carcasses detected after the last search numer $J$), all other values $\hat{\theta}_{(J-1)}^{(s,f)}, \hat{\theta}_{(J-2)}^{(s,f)}, ..., \hat{\theta}_{(j)}^{(s,f)}$ can be successively determined via the equation

$$\hat{\theta}_{(m)}^{(s,f)} = \frac{1}{\hat{s}_M(D_{(m,m-1)}, \vec{X}_i) \cdot \hat{f}(\vec{X}_i)} - K \cdot \sum_{n=m+1}^{J} \hat{\theta}_{(n)}^{(s,f)} L_j^{(n)}. \tag{8}$$

#### Decomposition and falling outside the search area

If decomposition time $\hat{t}_d(\vec{X}_i)$ is experimentally assessed independently of persistence probability, in the last iterative step (i.e., calculating $\hat{\theta}_{(j)}^{(s,f)}$ from equation 7), we set $L_j^{(n)} = 0$ if $D_{(j,n)} > \hat{t}_d(\vec{X}_i)$ holds. Thus, we incorporate the effect that carcasses can't be found after more than $\hat{t}_d(\vec{X}_i)$ days.

$\hat{\theta}_{(j)}^{(s,f)}(\vec{X}_i)$ now can be easily extended to incorporate the amount of carcasses dying outside the search area by simply considering

$$\frac{\hat{\theta}_{(j)}^{(s,f)}(\vec{X}_i)}{\hat{A}_{in}(\vec{X}_i)} =: \hat{\theta}_{(j)}(\vec{X}_i) \tag{9}$$

instead.

#### Imputation of missing values

It may arrive that certain (sub-)areas/sites can't be monitored during certain searches, e.g., due to extreme high vegetation or agricultural use. If we don't want to discard all data from the corresponding search, we have to estimate and include appearing missing values. In the following, we use the techniques of (multiple) data imputation (2, 6, 7). On the one hand, this approach offers an elegant method to impute data sensitively oriented on time- and space-dependent carcass numbers. On the other hand, multiple imputation techniques (including natural random spread around the expected value) fit well together with the resampling-method used for variance estimation (c.f., following section).

To impute missing values, we first sum up all corrected values (given by equation 9) for each existing date-site-combination. Subsequently, we fit a Generalized Additive Model (GAM) to the data, depending on the date (considered as a numerical variable) and the site (6). Here, we choose a GAM over a GLM(M) due to the high nonlinear time-dependent pattern, bird/bat migration usually shows. Finally, we impute the expected values $\hat{\theta}_1^{imput}, ..., \hat{\theta}_K^{imput}$ as predicted by the GAM. Since we deal with count data, we specify Poisson-distributed values $\hat{\theta}_k^{imput}$ within this regression model, which can be easily changed e.g. to a negative-binomial model if data show overdispersion (8, 9).

Finally, we obtain estimates $\hat{\theta}_1 ..., \hat{\theta}_I$ and imputed values $\hat{\theta}_1^{imput}, ..., \hat{\theta}_K^{imput}$ and define the final estimator of the sum of carcasses (for the period and area of interest) via

$$\hat{N} = \sum_{i=1}^{I} \hat{\theta}_i + \sum_{k=1}^{K} \hat{\theta}_k^{imput}. \tag{10}$$

### Monte-Carlo-based resampling-scheme

Here, we present and motivate the detailed resampling scheme in order to calculate confidence intervals in conjunction with non-parametric bootstrap methods.
We are interested in the average spread of $\hat{N}$ around the true value $N$. Thus the question is: If there are $N$ birds/bats, colliding with the structure of interest in the given area and period of interest. And if it would be possible to repeat the subsequent (random) processes of falling into the search area, removal by scavengers, detection by searchers, etc. several times – including the performance of additional experiments to approximate $f(), s(), t_d()$ and $A_{in}()$, as well as the final calculation of $\hat{N}$. How would the spread of $\hat{N}$ around $N$ qualitatively and quantitatively look like?
In praxis, $N$ is unknown and there is only one single realization $N^F$ from $N$ (and the thereupon calculated $\hat{N}$) available. However, we can virtually recreate these random processes by Monte-Carlo-simulations. The corresponding resampling mechanisms consists of the following three steps:

1. We use $\hat{N}$ as an approximation of $N$ and split it up into single observations.
2. Based on this, we virtually imitate all above mentioned random processes of removal, detection, etc., leading to a "virtual set of detected carcasses" $N_i^F$. Here, probabilities used in Monte-Carlo simulations are deduced from regression models of the corresponding experimental data
3. Finally, based on $N_i^F$, we calculate a new estimate $\hat{N}_i$ as described within the section "Deducing the estimator $\hat{N}$". Only differences are: we don't use the original experimental data to calculate correction terms, but calculate regression models based on (newly generated) resamples of the experimental data, mimicking a new performance of these experiments. Furthermore, for imputation, we artificially spread

the data around their expected value (based on regression standard errors as well as on the natural spread of the data).
